# Combinatorial Treatment Increases IKAP Levels in Human Cells Generated from Familial Dysautonomia Patients

**DOI:** 10.1101/524587

**Authors:** Sivan Yannai, Jonathan Zonszain, Maya Donyo, Gil Ast

**Author notes:** Address: Department of Human Molecular Genetics and Biochemistry, Sackler Faculty of Medicine, Tel Aviv University, Ramat Aviv 69978, Israel.

## Abstract

Familial Dysautonomia (FD) is an autosomal recessive congenital neuropathy that results from a point mutation at the 5’ splice site of intron 20 in the *IKBKAP* gene. This mutation decreases production of the IKAP protein, and treatments that increase the level of the full-length *IKBKAP* transcript are likely to be of therapeutic value. We previously found that phosphatidylserine (PS), an FDA-approved food supplement, elevates IKAP levels in cells generated from FD patients. Here we demonstrate that combined treatment of cells generated from FD patients with PS and kinetin or PS and the histone deacetylase inhibitor trichostatin A (TSA) resulted in an additive elevation of IKAP compared to each drug alone. This indicates that the compounds influence different pathways. We also found that pridopidine enhances production of IKAP in cells generated from FD patients. Pridopidine has an additive effect on IKAP levels when used in combination with kinetin or TSA, but not with PS; suggesting that PS and pridopidine influence *IKBKAP* levels through the same mechanism. Indeed, we demonstrate that the effect of PS and pridopidine is through sigma-1 receptor-mediated activation of the BDNF signaling pathway. A combination treatment with any of these drugs with different mechanisms has potential to benefit FD patients.

## Introduction

Familial Dysautonomia (FD) is an autosomal recessive congenital neuropathy that is characterized by abnormal development and progressive degeneration of the sensory and autonomic nervous systems ^1–3^. The gene associated with the disease is *IKBKAP*, which encodes a 150-kDa protein called IκB kinase complex-associated protein (IKAP) ^4, 5^. The mutation observed in 99.5% of FD patients is a transition from T to C at position 6 of the 5’ splice site of intron 20 ^1, 6^. This mutation occurs almost exclusively in the Ashkenazi Jewish population, with carrier frequency ranging from 1 in 32 to as high as 1 in 18 in those of Polish descent ^1, 7^. The mutation causes a shift in the splicing pattern of the *IKBKAP* pre-mRNA. Normally, exon 20 is constitutively included in the mature mRNA, but in the nervous systems of FD patients exon 20 is mainly skipped ^4, 8^. Interestingly, in non-nervous system tissues of FD patients, both wild-type (WT) and mutant *IKBKAP* mRNA are observed in varying ratios ^4^. We demonstrated previously that the affinity of the splicing factor U1 for the mutated 5’ splice site is reduced compared to that for the WT 5’ splice site ^9^.

The function of IKAP has been interrogated using both cellular and animal models; these models have also been important in analyses of potential therapeutic agents ^10^. Although IKAP is mostly localized in the cytoplasm, it was initially identified as subunit of the elongator complex which assists RNA polymerase II in transcription in the nucleus, affecting the transcript elongation of several genes ^6, 11–13^. IKAP is also implicated in regulation of the JNK signaling pathway ^14, 15^, tRNA modification ^16, 17^, cell adhesion, cell migration, and cytoskeleton stability and dynamics ^13, 18–20^. IKAP is also crucial for oligodendrocyte differentiation and/or myelin formation ^21, 22^, and vascular and neural development during embryogenesis ^19, 23, 24^.

Previous studies have shown that increasing the level of the full-length *IKBKAP* transcript is likely to be of therapeutic value. A number of strategies have been identified that increase the inclusion level of exon 20 in cells derived from FD patients or FD mouse models, and a platform has been developed to screen for potential small molecules that can affect *IKBKAP* splicing ^25–29^. We previously demonstrated that phosphatidylserine (PS) elevates *IKBKAP* transcription and, as consequence, IKAP protein levels in cells generated from FD patients (FD cells) and in humanized FD mice ^30, 31^. PS treatment releases FD cells from cell-cycle arrest ^30^, affects genes involved in Parkinson’s disease ^31^, and improves axonal transport ^20, 32^. PS treatment upregulates *IKBKAP* transcription by CREB and ELK1, which bind to the *IKBKAP* promoter region, activation of the mitogen-activated protein kinase (MAPK) pathway ^33^. PS has also been evaluated in a clinical trial in FD patients with positive results ^34^ Thus, fibroblasts generated from FD patients are a valid system for screening of potential drugs and therapies.

The main goal of this research is to explore therapeutic approaches which will improve the quality of life for FD patients, either by discovering new therapies or improving the effect of known ones. A good therapy for FD would therefore be one that either affects the transcription level or elevates the inclusion level of *IKBKAP*. We examined the combinations of PS with additional agents to achieve a synergistic affect. Kinetin, a plant cytokinin, was previously shown to increase the inclusion levels of exon 20 of *IKBKAP* in cells derived from FD patients; however, the effective dosage in FD patients led to severe side effects ^30, 39–41^. Thus, a low dose of kinetin combined with PS might be beneficial for FD patients. Inhibition of histone deacetylase (HDAC) leads to chromatin relaxation and promotes transcription of certain genes and inclusion of certain exons ^42, 43^. HDAC inhibitor trichostatin A (TSA) promote transcription by selectively inhibiting the class I and II mammalian histone deacetylase ^44, 45^. HDAC inhibitors have potential in treatment of neurodegenerative disorders as they play a crucial protective role in neurodegeneration ^46^. Here we show that combinations of PS either with kinetin or with TSA had additive effects on IKAP levels in FD cells. We also show that pridopidine, which is a dopaminergic stabilizer that has been evaluated as a treatment for Huntington’s disease ^35–38^, elevates *IKBKAP* transcription and as consequence IKAP protein levels. Combinations of pridopidine either with kinetin or with TSA had additive effects on IKAP levels in FD cells. However, PS and pridopidine did not have an additive effect on IKAP levels in FD cells, suggesting that these two compounds have the same mechanism of action. We provide evidence that PS activated the sigma-1 receptor (Sig-1R) in FD cells. This leads to activation of the MAPK signaling pathway by brain-derived neurotrophic factor (BDNF). Thus, the use of two drugs that act on different pathways have an additive effect on IKAP level and have potential for treatment of FD patients and possibly other disorders.

## Results

### Combined treatments with phosphatidylserine improve treatment effect observed compared to each drug alone

We evaluated several possible therapeutic agents used in combination for the effect on IKAP levels in FD cells. We examined combinations of either PS with kinetin, which acts on splicing, and then with TSA, which is an HDAC inhibitor. In FD cells, the combination of PS and kinetin led to elevation in IKAP protein level by 3.33 and 1.56 fold compared to PS and kinetin alone, respectively (Fig. 1A, ***p≤0.005 and *p≤0.05).

**Figure 1:**
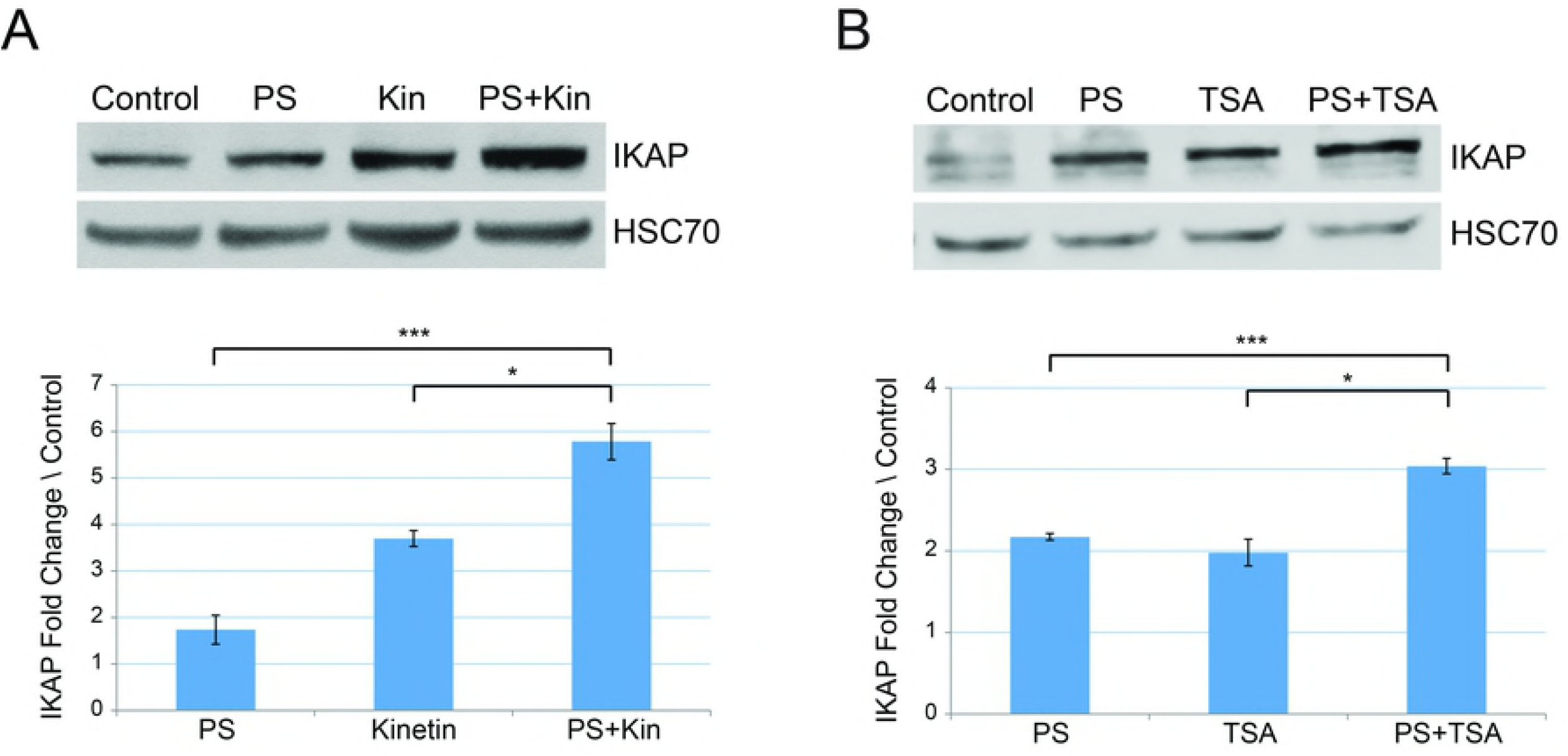
Combined treatments with phosphatidylserine elevates IKAP protein level more efficiently than either drug alone. FD cells were treated for 5 days either with **(A)** 50 μg/ml PS or 10 μM kinetin (Kin) or the combination of both drugs, and **(B)** 50 μg/ml PS or 100 ng/ml TSA or the combination of both drugs. Upper panels: Western blotting of FD cell lysates after indicated treatments. Proteins were extracted, and western blots were analyzed by using anti-IKAP antibody or HSC-70 antibody; the latter was used as a protein-loading control. Lower panels: Quantification of IKAP fold change levels normalized to HSC70 and relative to control (vehicle only). All quantifications were done using FusionCapt software. Asterisks denote statistically significant differences (*P ≤ 0.05 and ***P ≤ 0.005) relative to each control; Student’s *t*-test.

We then evaluated PS and TSA treatment and found that the combination resulted in the highest elevation of IKAP protein level, and IKAP levels were increased by 1.4 and 1.53 fold compared to PS and TSA alone, respectively (Fig. 1B, ***p≤0.005 and *p≤0.05). We thus found that several drugs can elevate IKAP protein level when used in combination compared to the effect of each drug alone.

### Pridopidine elevates IKAP protein levels in cells generated from FD patients

Pridopidine was developed for symptomatic treatment of Huntington’s disease ^37^, which, like FD, is a neurodegenerative disorder. In order to investigate whether pridopidine has an effect on *IKBKAP* transcription, FD cells were treated with concentrations of pridopidine ranging from 0–10,000 nM. We observed the largest increase in IKAP protein and *IKBKAP* transcript levels at 500 nM of pridopidine. The addition of 500 nM pridopidine increased *IKBKAP* expression levels by 1.43-fold after 5 days of treatment and the amount of IKAP protein by 4-fold after 10 days of treatment relative to untreated FD cells (Fig. 2A and 2B, *p≤0.05 and ***p≤0.005). Pridopidine did not alter the ratio of the isoform that included exon 20 (WT) relative to the isoform in which exon 20 is skipped (Mut) but rather elevated the total amount of both isoforms. This suggests that, like PS ^30^, pridopidine increases IKAP levels by elevating *IKBKAP* transcription level rather than by affecting the inclusion level of exon 20 (Fig. S1).

**Figure 2:**
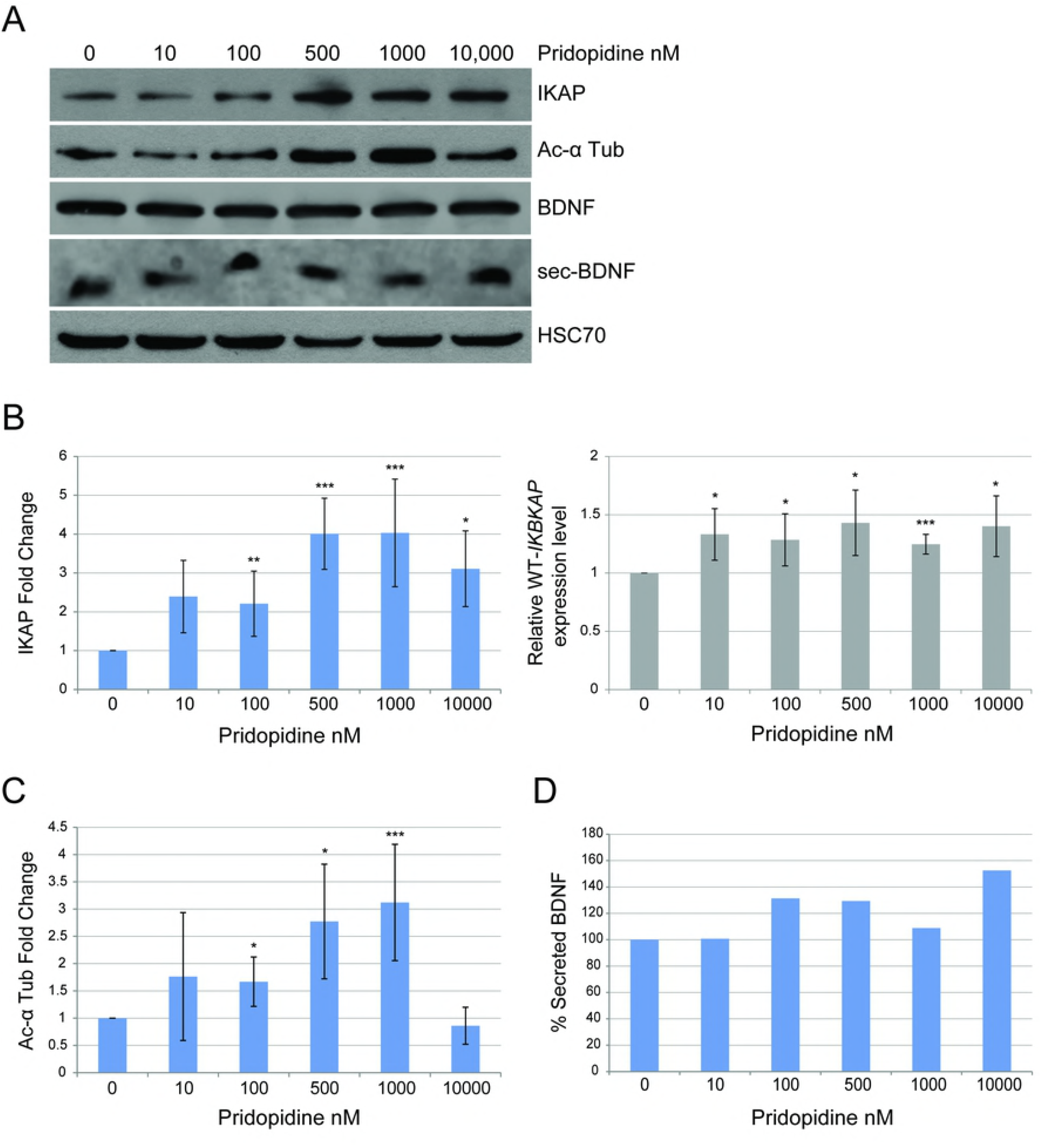
Pridopidine elevates IKAP protein level and affects acetylated α-tubulin levels and BDNF secretion in FD cells. **(A-D)** FD cells were treated with 0, 10, 100, 500, 1000, and 10,000 nM of pridopidine. Control treatment was done with vehicle only. **(A)** Western blotting of FD cell lysates with and without pridopidine treatment for 10 days. Blot was probed using anti-IKAP, anti-acetylated α-tubulin, anti-BDNF, and anti-HSC-70 antibodies. HSC-70 was analyzed as a protein-loading control. **(B)** Left panel: Fold change levels of IKAP relative to control analyzed after a 10-day treatment; levels were normalized to HSC70. Right panel: *IKBKAP* expression levels relative to control analyzed by qRT-PCR from RNA extracted from FD cells after 5 days of treatment. **(C)** Fold change levels of acetylated α-tubulin relative to control normalized to HSC70 analyzed after 10 days of treatment. **(D)** Quantification of percent BDNF secretion after a 72-hour incubation with pridopidine. All quantifications were done using FusionCapt software. Asterisks denote statistically significant differences (*P ≤ 0.05, **P ≤ 0.01, and ***P ≤ 0.005) relative to control (vehicle only); Student’s *t*-test.

Moreover, as does PS ^32^, pridopidine affects the levels of acetylated α-tubulin with the highest effect of 3.12-fold elevation observed at 1000 nM concentration (Fig. 2A and 2C, ***p≤0.005). Since pridopidine was shown to affect BDNF secretion ^37^, we examined this effect in FD cells by incubating the cells with a series of concentrations of pridopidine for 72 hours, and then we analyzed the medium for BDNF. We demonstrate that pridopidine affected BDNF in FD cells both by elevating the amount of BDNF secreted compared to levels secreted by untreated cells (Fig. 2A and 2D), and by enhancing the expression of several BDNF-induced genes (Fig. S2).

### Combined treatments with pridopidine improve treatment effect observed compared to each drug alone

We also examined combinations of either pridopidine with kinetin and TSA. In FD cells, pridopidine combined with kinetin led to elevation in IKAP protein level by 3.31 and 1.26 fold compared to pridopidine and kinetin alone, respectively (Fig. 3A, ***p≤0.005 and *p≤0.05). Pridopidine combined with TSA elevated IKAP levels by 1.19 and 1.36 fold compared to pridopidine and TSA alone, respectively (Fig. 3B, *p≤0.05). This indicates that kinetin, like PS, can elevate IKAP protein level when used in combination compared to the effect of each drug alone.

**Figure 3:**
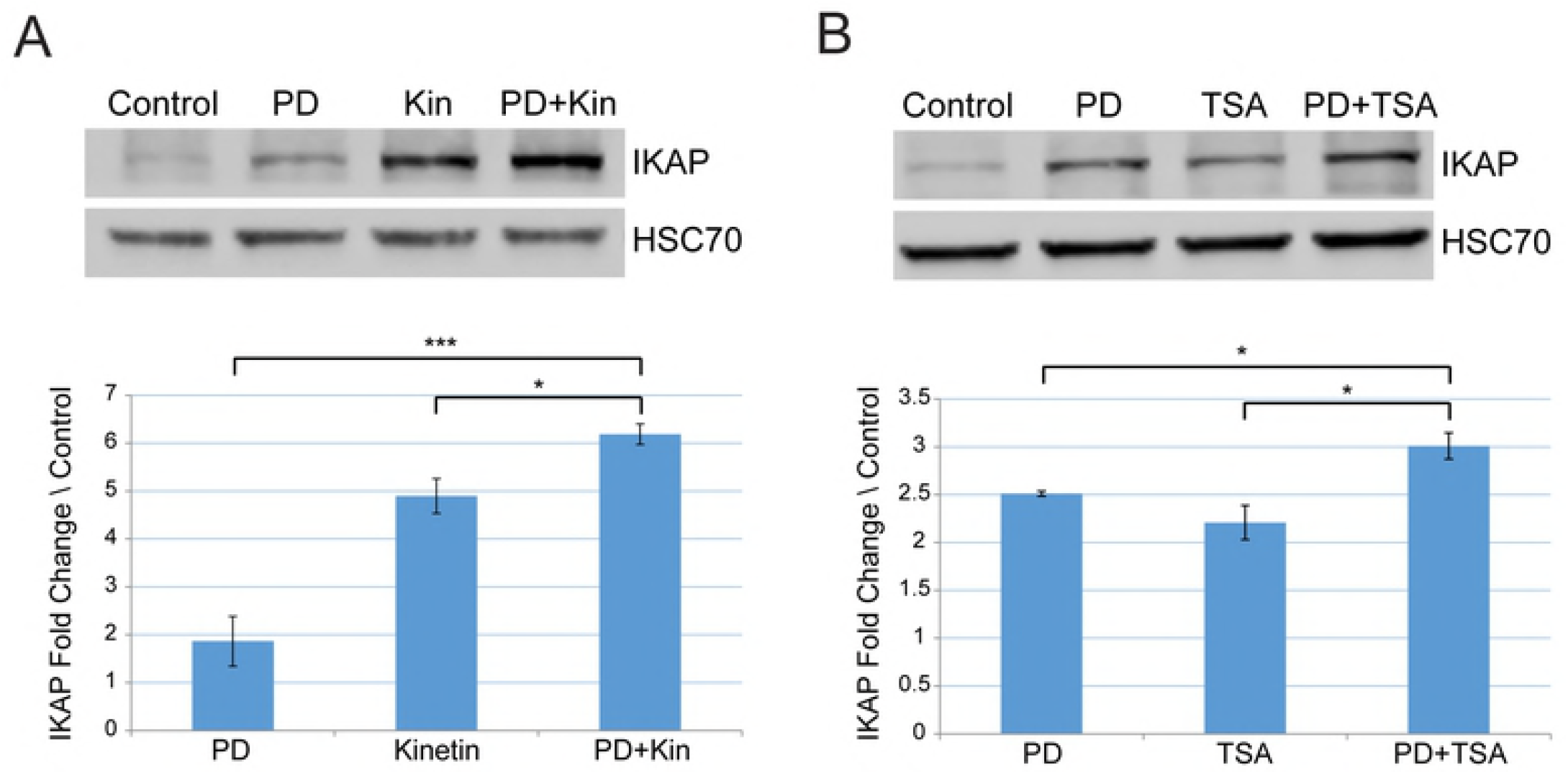
Combined treatments with pridopidine elevates IKAP protein level more efficiently than either drug alone. Western blotting of FD cell lysates after indicated treatments. Proteins were extracted, and western blots were analyzed by using anti-IKAP antibody or HSC-70 antibody; the latter was used as a protein-loading control. FD cells were treated for 7 days either with **(A)** 500 nM pridopidine (PD) or 10 μM kinetin or and the combination of both drugs, and **(B)** 500 nM pridopidine or 100 ng/ml TSA or the combination of both drugs. Upper panels: Western blotting of FD cell lysates after indicated treatments. Proteins were extracted, and western blots were analyzed by using anti-IKAP antibody or HSC70 antibody; the latter was used as a protein-loading control. Lower panels: Quantification of IKAP fold change levels normalized to HSC70 and relative to control (vehicle only). All quantifications were done using FusionCapt software. Asterisks denote statistically significant differences (*P ≤ 0.05 and ***P ≤ 0.005) relative to each control; Student’s *t*-test.

### Phosphatidylserine activates MAPK signaling pathway through sigma-1 receptor in FD cells

We recently showed that PS elevates *IKBKAP* transcription level through activation of MAPK pathway and found that inhibition of MEK1 and MEK2 decreased *IKBKAP* expression ^33^. The effects of pridopidine on expression of genes involved in the BDNF signaling pathway are mediated through the Sig-1R receptor ^37, 38^. We thus showed that the combination of PS and pridopidine did not lead to higher levels of IKAP compared to treatment with either drug alone (Fig. 4A). This was not unexpected given that both act on the MAPK pathway. In order to examine whether PS activates MAPK through Sig-1R, FD cells were treated either with PS or pridopidine, with or without NE-100, which is a Sig-1R antagonist. Pretreatment with NE-100 resulted in decreased IKAP levels in cells treated with PS than in cells only treated with PS (Fig. 4B, **p≤0.01). This indicates that elevation of IKAP levels upon PS treatment is mediated by Sig-1R. Although pridopidine activity was shown to be mediated through Sig-1R ^37, 38^, pretreatment with NE-100 did not alter the levels of IKAP in pridopidine-treated FD cells (Fig 4B). In order to further investigate the effect on *IKBKAP* we then examined MAPK/ERK inhibitor U0126 treatment combined with pridopidine. Pretreatment with U0126 did reduce efficacy of pridopidine treatment (Fig. 4C, *p≤0.05), demonstrating that pridopidine influences *IKBKAP* transcription through the MAPK/ERK signaling pathway. These results indicate that pridopidine appears to mediate signal transduction that enhances IKAP production through other cellular receptors in addition to Sig-1R.

**Figure 4:**
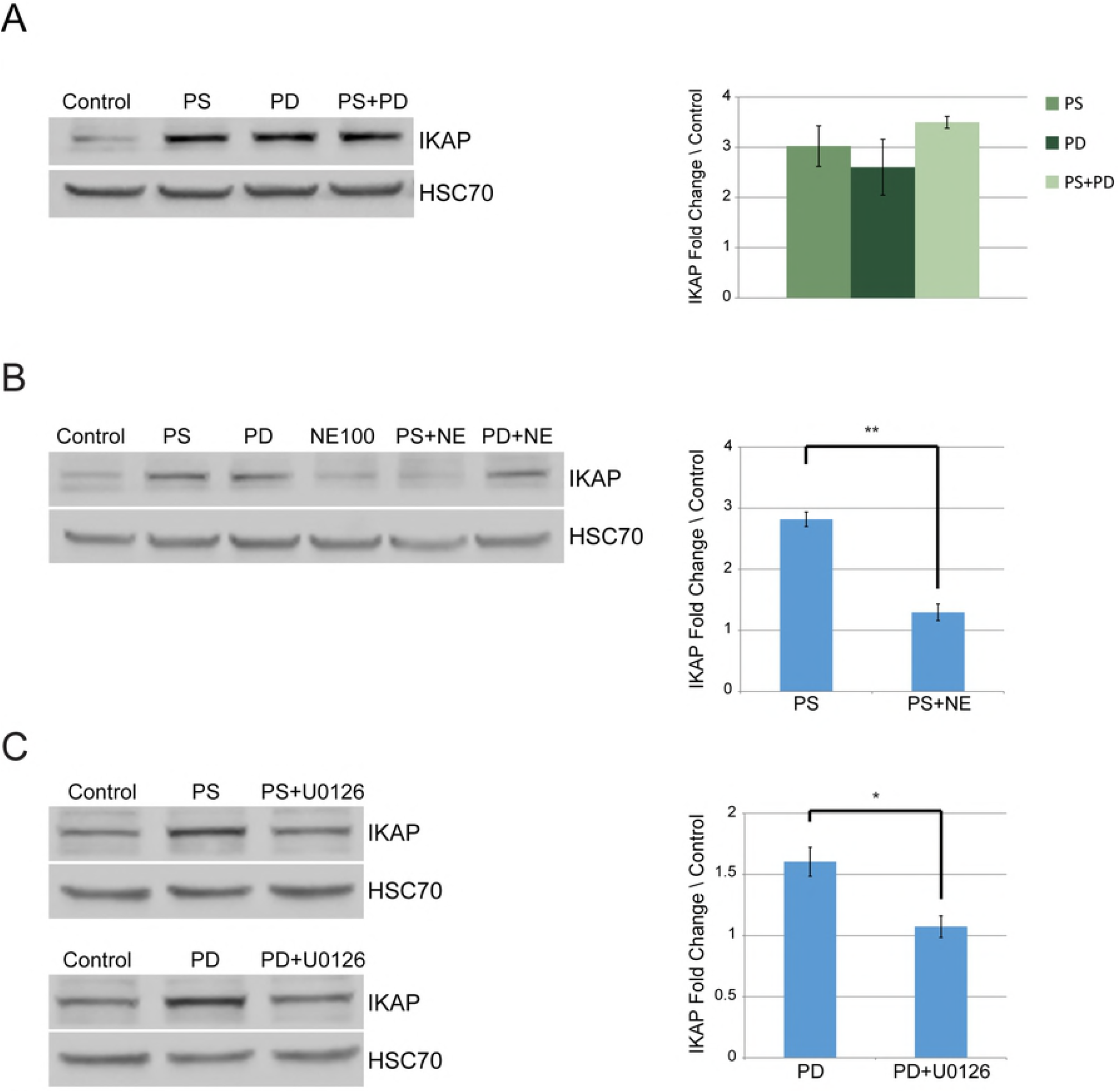
Phosphatidylserine and pridopidine increase IKAP levels in FD cells through the same mechanism. **(A)** FD cells were treated either with 50 μg/ml of PS or 500 nM of pridopidine or the combination of both drugs. Proteins were extracted and analyzed after 7 days. Left panel: Western blot of extracted proteins analyzed using anti-IKAP antibody. Right panel: IKAP fold change levels normalized to HSC70 and relative to control (vehicle only). **(B)** Left panel: Western blotting with anti-IKAP antibody of lysates of FD cells treated with 50 μg/ml PS, 500 nM of pridopidine, 2 μM sigma-1 receptor inhibitor NE-100, or indicated combinations. Right panel: IKAP fold change levels normalized to HSC70 and relative to control. **(C)** Left panel: Western blotting using anti-IKAP antibody of lysates of FD cells treated with 50 μg/ml PS or 500 nM pridopidine with or without 2 μM MAPK inhibitor U0126. Right panel: IKAP fold change levels normalized to HSC70 and relative to control. All quantifications were done using FusionCapt software. Asterisks denote statistically significant differences (*P ≤ 0.05 and **P ≤ 0.01) relative to control (vehicle only treated cells); Student’s *t*-test.

Since we demonstrated that the effect of PS on *IKBKAP* involves Sig-1R and MAPK signaling activation, we asked whether this activation is mediated through BDNF. In order to examine BDNF involvement in response to PS treatment we evaluated expression of several BDNF-induced genes in FD cells. PS treatment enhanced the expression of known BDNF targets including *CDKN1A, HOMER1, RASSF8, EGR1, SRXN1, RGS17*, and *BAIAP2* (Fig. 5). Thus, PS affects *IKBKAP* transcription level by activating sigma-1 receptor, which leads to activation of the BDNF signaling pathway.

**Figure 5:**
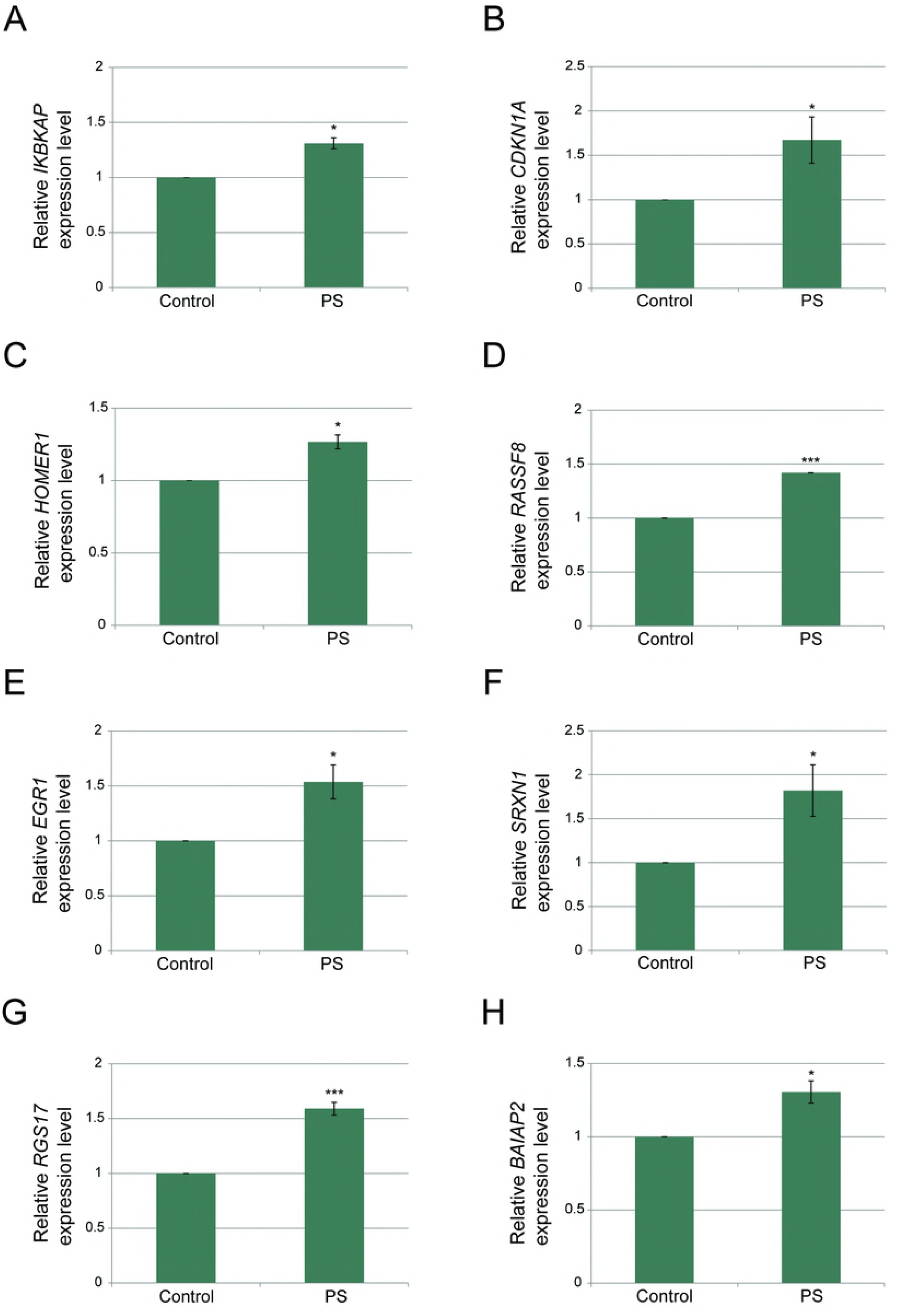
Phosphatidylserine activates the BDNF-mediated signaling pathway. **(A-H)** FD cells were treated with 50 μg/ml PS or a vehicle only control, and RNA was extracted 5 days after the treatment. qPCR was used to quantify **(A)** WT *IKBKAP*, **(B)** *CDKN1A*, **(C)** *HOMER1*, **(D)** *RASSF8*, **(E)** *EGR1*, **(F)** *SRXN1*, **(G)** *RGS17*, and **(H)** *BAIAP2* mRNA transcripts. All values were normalized to *LZIC*, which did not change as a result of PS treatment. Asterisks denote statistically significant differences (*P ≤ 0.05 and **P ≤ 0.01) relative to control (vehicle only); Student’s *t*-test.

## Discussion

The main purpose of this study was to identify potential new therapies, and improve the therapeutic effect on IKAP in a way that could be beneficial for FD patients. In our pursuit to find a way to improve the therapeutic effect in FD, we considered different agents with the potential to affect *IKBKAP*. Several potential therapies for FD have been investigated including PS ^30–32, 34^, kinetin ^40, 41^, tocotrienols ^47, 48^, and the green tea component epigallocatechin gallate^49^. Other therapy strategies are based on the FD mutation acting by altering gene splicing in the nerve system in a tissue-specific manner ^25, 27, 50–52^. We focused on PS since it is a well-studied, FDA-approved food supplement that upregulates IKAP production and has no known side effects; therefore, we sought to identify other compounds that could be used in combination with PS to benefit patients. Pridopidine was originally developed for symptomatic treatment of Huntington’s disease ^37^, and here we showed that pridopidine elevates IKAP levels in FD cells. PS was tested in combination with pridopidine and with several other drugs that have the potential to increase IKAP production through different mechanisms. We demonstrated how the combination of either PS or pridopidine with a low concentration of kinetin had an additive effect on IKAP in FD cells. Although kinetin effectively increases IKAP levels in FD cells, the drug causes severe side effects ^30, 39–41^. Our data indicates that use of kinetin, even at low concentrations, in combination with PS or pridopidine warrants further testing.

Inhibition of HDACs has been previously linked to anticancer effects ^53, 54^, and recent studies have demonstrated the synergistic effect of HDAC inhibitors in combination with standard chemotherapy for treatment of cancer ^55, 56^. We previously showed that in dorsal root ganglia, deficiency in IKAP results in elevation of histone deacetylase HDAC6 and reduction in the level of acetylated α-tubulin. PS acts as an inhibitor of HDAC6, a class II HDAC, resulting in elevated α-tubulin levels and enhanced nerve growth factor movement along the microtubules ^20, 32^. As PS is an inhibitor of a class II HDAC, we tested TSA, a class I inhibitor in combination with PS and pridopidine. Additive effects were observed in both cases, demonstrating the therapeutic potential of combining treatments with different mechanisms.

Finally, we demonstrated that in cells treated with the combination of PS and pridopidine, the increase in IKAP production was similar to cells treated with either drug alone. In contrast, the combinations of PS or pridopidine with either kinetin or TSA did have additive effects on the level of the IKAP protein. This was not unexpected given that both PS and pridopidine act on the MAPK pathway. This led us to investigate more deeply the mechanisms of action of PS and pridopidine. Treatment with PS was recently shown to lead to activation of the MAPK signaling pathway. Downstream regulators of MAPK signaling CREB and ELK1 bind to the *IKBKAP* promoter to upregulate its transcription ^33^. However, how PS ignites this cascade was not known. Here we showed that pretreatment with NE-100 resulted in decreased IKAP levels following PS treatment. This indicates that PS affects *IKBKAP* through the activation of sigma-1 receptor. Sig-1R is highly expressed in cells of the central nervous system and is located in the endoplasmic reticulum membrane ^57, 58^.

Sigma receptors have emerged as targets for novel therapeutic applications in neurodegenerative diseases ^59, 60^, since their activation is linked with neuroprotection ^61^. Sigma-1 receptor activation results in expression of BDNF, which mediates phosphorylation of the tyrosine kinase B (TrkB) ^62^, which initiates a signaling cascade that involving MEK and ERK that causes activation of MAPK ^63, 64^. We demonstrated that PS treatment of cells activated a BDNF-mediated activation of MAPK signaling pathway by examining the effect of PS treatment on several genes known to be induced by BDNF. Moreover, the Sig-1R inhibitor NE-100 blocked upregulation of IKAP caused by PS treatment. The fact that pre-treated FD cells show lower BDNF activity is consistent with the previous finding that TrkB and BDNF are crucial during sympathetic nervous system development ^65^. Although pridopidine activity was shown to be mediated through Sig-1R ^37, 38^, pretreatment with the Sig-1R inhibitor NE-100 did not alter the levels of IKAP in pridopidine-treated FD cells. Pretreatment with MAPK/ERK inhibitor U0126 did reduce efficacy of pridopidine treatment. Thus, the influence of pridopidine on IKAP production might involve other cellular receptors that activate the MAPK pathway. That PS and pridopidine do not have additive effects on IKAP production in FD cells likely results from their effects BDNF activity.

In conclusion, the notion that different neurodegenerative disorders have common mechanisms suggested to us that a therapy that has proven effective in treatment of Huntington’s disease might have activity in FD. This proved to be the case for pridopidine, which increased production of IKAP in FD cells. Moreover, drugs that act through different mechanism can be combined to yield an additive effect; further testing is warranted to determine whether any of the combinations tested here are synergistic. In addition understanding the regulation of *IKBKAP* is also important and can help identify new strategies for therapy. Therefore, the results in this study have great value and not only for FD patients.

## Materials and methods

### Cell culture

Human FD fibroblast cells were obtained from the appendices of FD patients and immortalized using telomerase activation ^30^. The human FD cell line was cultured in Dulbecco’s modified Eagle’s medium, supplemented with 4.5 g/ml glucose, 2 mM L-glutamine, 100 U/ml penicillin, 0.1 mg/ml streptomycin, and 20% fetal calf serum. Cells were grown in a 10-cm culture dish, under standard conditions, at materials were purchased from Biological Industries. The cells were seeded and on the next day drug or drug combinations were added. Every two days the medium was replaced and fresh treatments were added. The cells were split as needed during the treatment period to allow proper growth.

### PS treatment

InCog™, a lipid composition containing PS-omega 3, DHA enriched, referred to here as PS, was dissolved in organic solvent medium chain triglycerides (MCT). Both PS and MCT were obtained from Enzymotec. In all treatments PS was used at 50 μg/ml and was compared to its solvent as a control.

### Pridopidine treatment

Pridopidine was obtained by Teva Pharmaceutical Industries Ltd. Followed by a calibration, in all treatments Pridopidine was used at 500 nM and was compared to its solvent as a control. Pridopidine was dissolved in doubly-distilled water (DDW).

### Kinetin treatment

Kinetin was kindly provided by David Brenner and was dissolved in DMSO, and then diluted in fresh medium (0.1% DMSO). Followed by a calibration, in all treatments kinetin was used at 10 nM and was compared to its solvent only as control.

### TSA treatment

TSA was purchased from Sigma and dissolved in DMSO according to manufacturer’s instructions, and then diluted in DDW (0.1% DMSO). TSA was used at 100 ng/ml and compared to its solvent only as a control.

### Inhibitor treatments

U0126 was purchased from Calbiochem and dissolved in DMSO according to the manufacturer’s instructions. NE-100 was kindly provided by Teva and was dissolved in DDW. U0126 and NE-100 were used at 2 μM. For all experiments in which these inhibitors were used, the inhibitor was added 1 hour prior to addition of the other drug.

### Protein purification and western blot

Total proteins were extracted from the cells using a hypotonic lysis buffer (50 mM Tris-HCl, pH 7.5, 1% NP40, 150 mM NaCl, 0.1% SDS, 0.5% deoxycholic acid, 1 mM EDTA) containing protease inhibitor and phosphatase inhibitor cocktails I and II (Sigma). After 20-min centrifugation at 14,000 g at 4 °C, the supernatant was collected and protein concentrations were measured using BioRad Protein Assay (BioRad). Secreted proteins were collected after a 72-h incubation by Amicon Ultra-15 Centrifugal Filter Units (Merck) according to manufacturer’s protocol. Proteins were separated by 10% or 12% SDS-PAGE and then electroblotted onto a Protran nitrocellulose transfer membrane (Schleicher & Schuell). Immunoblots were incubated with primary and secondary antibodies, and signal was enhanced using chemiluminescence (SuperSignal West Pico chemiluminescent substrate; Thermo Scientific). The signal was detected by exposure to X-ray film or using the Fusion FX7 image acquisition system (Vilber Lourmat). Data were quantified using ImageJ ^66^, or using the FusionCapt software. Reported are data from at least three separate experiments.

### Antibodies

Primary antibodies used for immunoblotting were as follows: anti-IKAP (Anaspec, cat# 54494), anti-acetylated α-tubulin (Sigma, cat# t7451), anti-BDNF (Alomone Labs, cat# ANT010), and anti-Hsc70 (Santa Cruz Biotechnology, cat# sc-7298). Secondary antibodies were donkey anti-rabbit IgG HRP (Abcam, cat# ab97064), or donkey anti-mouse IgG HRP (Abcam, cat# ab98799), as appropriate.

### RNA purification and quantitative RT–PCR

RNA was extracted from FD cells using TRI reagent (Sigma) and reverse transcribed using the SuperScript III First Strand kit (Invitrogen) with an oligo(dT) reverse primer. A qPCR analysis of mRNA expression from FD cells samples were conducted using KAPA SYBR Fast qPCR master mix (Kapa Biosystems) in a StepOne plus thermocycler PCR machine (Applied Biosystems) according to the manufacturer’s instructions. *LZIC* was used as endogenous control. The primer sequences are listed in Table 1S.

## Acknowledgments

This work was performed in partial fulfillment of the requirements for a Ph.D. degree of S.Y. at the Sackler Faculty of Medicine, Tel Aviv University. We are grateful for the support of the Israeli National Network of Excellence from Teva Pharmaceutical Industries Ltd. We are also grateful for Enzymotec for supplying PS. This paper is dedicated to the memory of David Brenner, former president of the Dysautonomia Foundation.

## Abbreviations

FD: Familial Dysautonomia
IKAP: IκB kinase complex-associated protein
MAPK: mitogen-activated protein kinase
BDNF: brain-derived neurotrophic factor
PS: phosphatidylserine
HDAC: histone deacetylase
TSA: trichostatin A
WT: wild type

